# Segmenting Microarrays with Deep Neural Networks

**DOI:** 10.1101/020404

**Authors:** Andrew Jones

## Abstract

Microarray images consist of thousands of spots, e ach of which corresponds to a different biological material. The microarray segmentation problem is to work out which pixels belong to which spots, even in presence of noise and corruption. We propose a solution based on deep neural networks, which achieves excellent results both on simulated and experimental data. We have made the source code for our solution available on Github under a permissive license.

## 1 Introduction

In a typical two-channel comparative hybridization experiment, RNA from a control sample is tagged with a green fluorescent dye while RNA from an experimental sample is tagged with a red fluorescent dye. The tagged samples are mixed, then hybridized to a microarray. A microarray is a small glass slide printed with a rectangular array of thousands of spots. Each spot contains a specific DNA probe sequence, which during hybridization can bind to complementary RNA from the sample. If a specific RNA sequence is expressed more strongly in the experimental sample than in the control sample, then the corresponding spot will fluoresce more strongly under a green laser than under a red. The difference in fluorescence can be measured by scanning the array with both kinds of laser, then comparing the scanned images using software.

To measure the expression difference of a DNA probe, the software needs to know which pixels belong to the spot corresponding to that probe and which pixels belong to the background. This is the microarray image segmentation problem. It is complicated by issues in the manufacture and processing of arrays, which leads variation in the location, shape, size and intensity of spots. The same issues can also corrupt whole regions with streaks or blotches as in Figure 1 within which the intensity of a pixel in a channel is not a reliable indicator of the corresponding sequence’s expression level.

**Figure 1:**
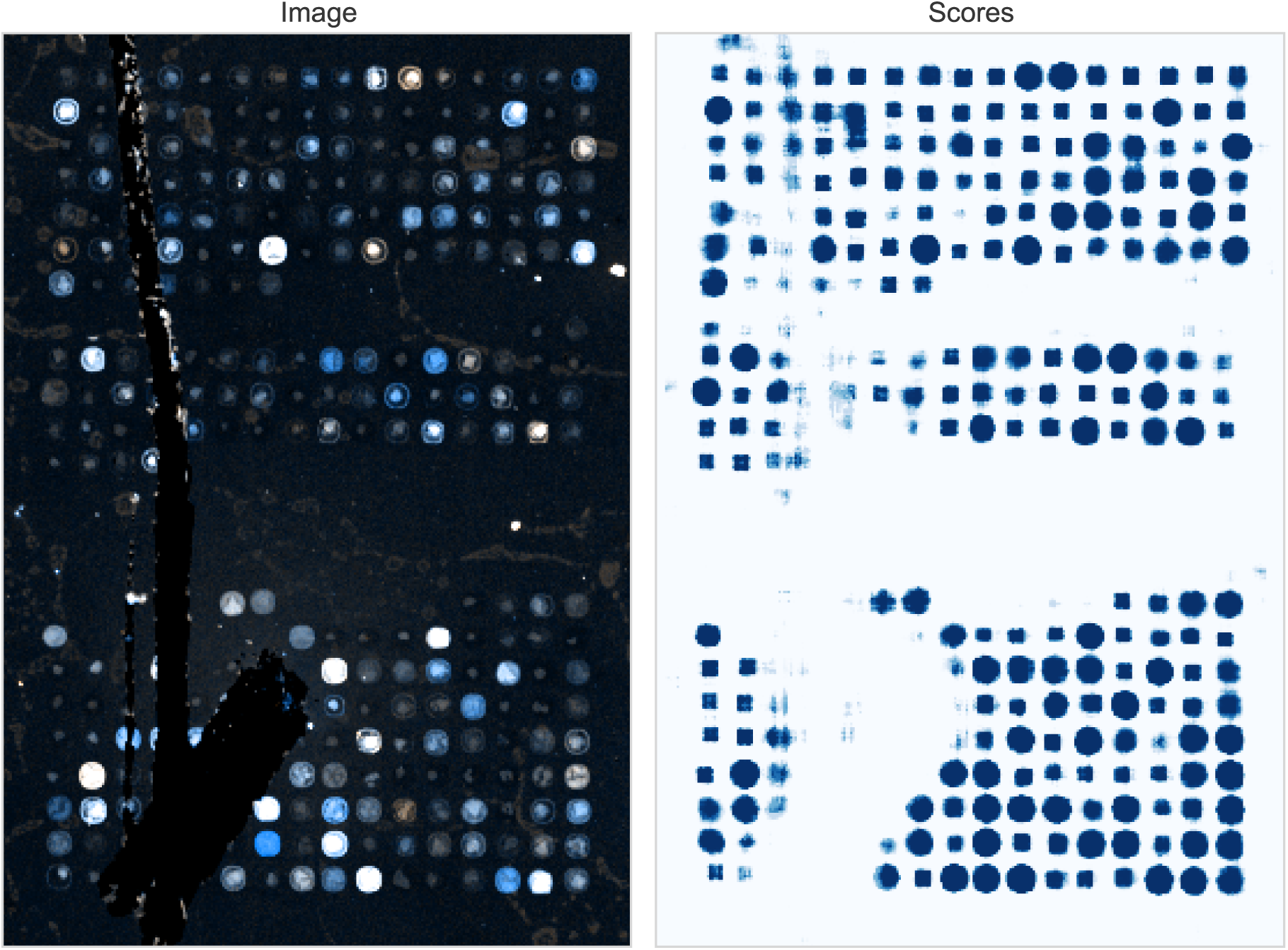
An example of a corrupted area in a sub-array, together with the scores reported by our neural network.

A variety of algorithms and software packages have been developed to solve the segmentation problem [1,5]. These algorithms and packages generally reflect approaches that have been applied to the more general problem of image segmentation [22], in which an image is broken down into regions corresponding to perceptually distinct objects or areas. One of the most successful approaches to image segmentation in recent years has been deep neural networks [16, 6, 18, 9].

### 1.1 Deep neural networks

For a detailed introduction to modern neural networks, see Bengio et al.’s upcoming book [2].

At a conceptual level, a deep convolutional neural network is formed from a stack of layers, with the input at the bottom and the output at the top. Each layer is composed of a large number of neurons, which look for specific patterns in the output of the layer below, and pass the intensity of the pattern to the layer above.

The neural network is taught which patterns to look for by reinforcement learning. In reinforcement learning, the network is repeatedly presented with samples from a training set. If the network generates the correct output, then the neurons that contributed to the output have their importance in the network increased. If the network gives an incorrect response, those same neurons have their importance reduced. Over thousands of samples, the network learns to generate the responses the trainer is looking for.

Neural networks have been an active area of research since the 1960s, but have undergone a renaissance in recent years thanks to significant improvements in both theory and hardware. The most important advancement has been the discovery of initialization and training schemes that work on networks with many layers. These ‘deep’ neural networks have produced impressive results in a wide range of computer vision problems.

Neural networks have been applied to the microarray image segmentation problem previously [19], but the networks used were shallow (three layers) and applied only as part of a larger heuristic. In contrast, our inspiration was Ciresan et al. [4], where a deep network was used to identify neuron membranes in electron microscopy images.

## 2 Methods

Our approach is similar to Ciresan et al.’s. To decide whether a pixel belongs to a spot or to the background, we take various windows centred on that pixel and ask a neural network to score each window on a scale of [0, 1]. A score near 0 means the network thinks the window is centred on a background pixel, while a score near 1 means it’s centred on a spot pixel.

Given the scores of all the windows centred on a pixel, the score of a pixel is their median. This leads to a score map similar to Figure 1, which can be thresholded to decide which pixels should be considered spot-pixels and which should be considered background.

### 2.1 Datasets

Our data was taken from Lehmussola et al.’s [15] evaluation of various approaches to the microarray segmentation problem. Although old, the data Lehmussola et al. used for their evaluations is freely available on the website associated with their paper, and several authors since have used it as a benchmark.

Lehmussola et al.’s benchmark uses two datasets. The first dataset is of simulated microarray images, created using the image generator of Nykter et al [17]. It contains 50 ‘high-quality’ images and 50 ‘low-quality’ images. The low quality set has a larger fraction of damaged or misplaced spots, and a higher level of background noise. Each image contains 1, 000 spots, and comes with a ground-truth channel that indicates which pixels belong to spots and which belong to the background.

The ground truth is a real-valued number [0, 1], but for the purposes of training and testing a classifier we assumed pixels with a ground truth less than 0.5 were background, while the rest belonged to spots.

The second dataset contains five real microarray images, all replicates from the same experiment. Since no groundtruth was available, we hand-labelled 3,000 spots in six blocks in one of the benchmark images. Each pixel was labelled as belonging to a good spot, belonging to a damaged spot, belonging to a missing spot, or left unlabelled to indicate that it belonged to the background. Our labellings were approximate rather than exact, as we believe that a large number of spots with mostly-correct labels form a better training set than a small number of spots with entirely correct labels.

As well as marking individual spots, we also used the image editor to outline blocks of spots, even if the spots within weren’t individually labelled. This let us sample random background pixels without accidentally choosing a spot pixel.

### 2.2 Preprocessing

To deal with the huge range of microarray pixel intensities, we used the log-intensity of the pixels, and normalized the logged images so that they had mean 0 and variance 1.

For the experimental benchmark, we generated windows of 61× 61 pixels, while for the simulated benchmark we generate windows of 41× 41 pixels. These each contained about 3 × 3 spots, with the spots in the simulated benchmark being smaller than those in the experimental benchmark. The windows have an odd width so that reflections and 90° rotations of the window would still be centred on the same pixel.

For each pixel, we generated 8 windows: the original window, its reflection, the three 90° rotations of the original window and the three 90° rotations of the reflected window. Each of these windows was scored by the network, and the median of the results was taken to be the score of the pixel.

### 2.3 Network architecture

Our neural network architecture is detailed in Table 2.3. It was derived from the architecture used by Ciresan et al, but with some differences.

First, our network did not use foveation or non-uniform sampling. We felt these would be counter-productive for our specific segmentation problem, as exact knowledge of where a spot’s neighbours are seems important in deducing where the spot itself is.

Second, our network used one fewer convolution layer, as the 41 × 41 windows used for the simulated data benchmark are smaller than the 64 × 64 windows used by Ciresan et al. With a 41 × 41 input, the output after three convolution layers and three max-pool layers had shape 1× 1; a fourth convolution layer would have added little.

Third, our network used several recent advances in neural network theory. These include overlapping convolution kernels [14], dropout layers [11] and ReLU nonlinearities [8]. Our network also used a smaller stride for its convolution layers, made possible by advances in GPGPU hardware.

**Table 1:** Network architecture

| Type | Channels | Kernel | Stride |
| --- | --- | --- | --- |
| Data | 2 |  |  |
| Convolution | 32 | $4 \times 4$ | 1 |
| ReLU | 32 |  |  |
| Max Pool | 32 | $4 \times 4$ | 2 |
| Convolution | 64 | $4 \times 4$ | 1 |
| ReLU | 64 |  |  |
| Max Pool | 64 | $4 \times 4$ | 2 |
| Convolution | 96 | $4 \times 4$ | 1 |
| ReLU | 96 |  |  |
| Max Pool | 96 | $4 \times 4$ | 2 |
| Fully-connected | 192 |  |  |
| ReLU | 192 |  |  |
| Dropout | 192 |  |  |
| Fully-connected | 2 |  |  |
| Sigmoid | 2 |  |  |

### 2.4 Network training

We trained two networks, one for each benchmark. This was necessary because of differences in array properties between the two benchmarks (like spots in the simulated benchmark being much smaller than those in the experimental benchmark).

In each case, the network was implemented using the Caffe framework [12]. The weights were initialized using the PReLU scheme [10], and the network was trained using SGD with momentum, weight decay and a stepped learning rate. For further details, the code and the model definitions can be found on Github.

#### 2.4.1 Simulated Data Training and Test Sets

We used low-quality images 1 through 25 from Lehmussola et al.’s simulated dataset to generate the training and test sets. In total, 800,000 windows were created, of which

- 25% were centred on pixels that belong to a spot.
- 25% were centred on pixels that belong to ‘damaged’ spots, which are either misshapen, an unusual size or misaligned.
- 12.5% were centred on pixels that belong to the background.
- 12.5% were centred on pixels that belong to the background near damaged spots.
- 12.5% were centred on pixels that belong to the background near the perimeter of a block of spots.
- 12.5% were centred on pixels that belong to the background between different spots.

The first two classes constituted the positive examples and were labelled 1, while the others were labelled 0. We originally sampled just from background pixels and spot pixels rather than from these specific classes, but found that this lead to an implicit ‘class imbalance’ problem. Because most spots are undamaged, the network wouldn’t learn to recognise damaged spots, and would instead predict pixels around damaged areas as if they were undamaged. Similarly, because most spots lie in the interior of a block of spots, the network wouldn’t learn to identify the boundary of a block and instead would continue to predict spots where none existed.

#### 2.4.2 Experimental Data Training and Test Sets

For the experimental benchmark we generated 400,000 windows in much the same way as for the simulated data, but using our hand-labellings as the ground truth. The reduced number of windows was because the number of hand-labelled spot pixels is far smaller than the number of spot pixels in the simulated data’s ground truths.

### 2.5 Visualization

Although it didn’t contribute directly to our results, we feel it’s worth discussing the visualization of two-channel microarray images. The most common way to visualize a two-channel microarray is to map the raw intensity values onto [0,255] and interpret them as the red and green channels of a 24-bit image, as on the left-hand side of Figure 2.

**Figure 2:**
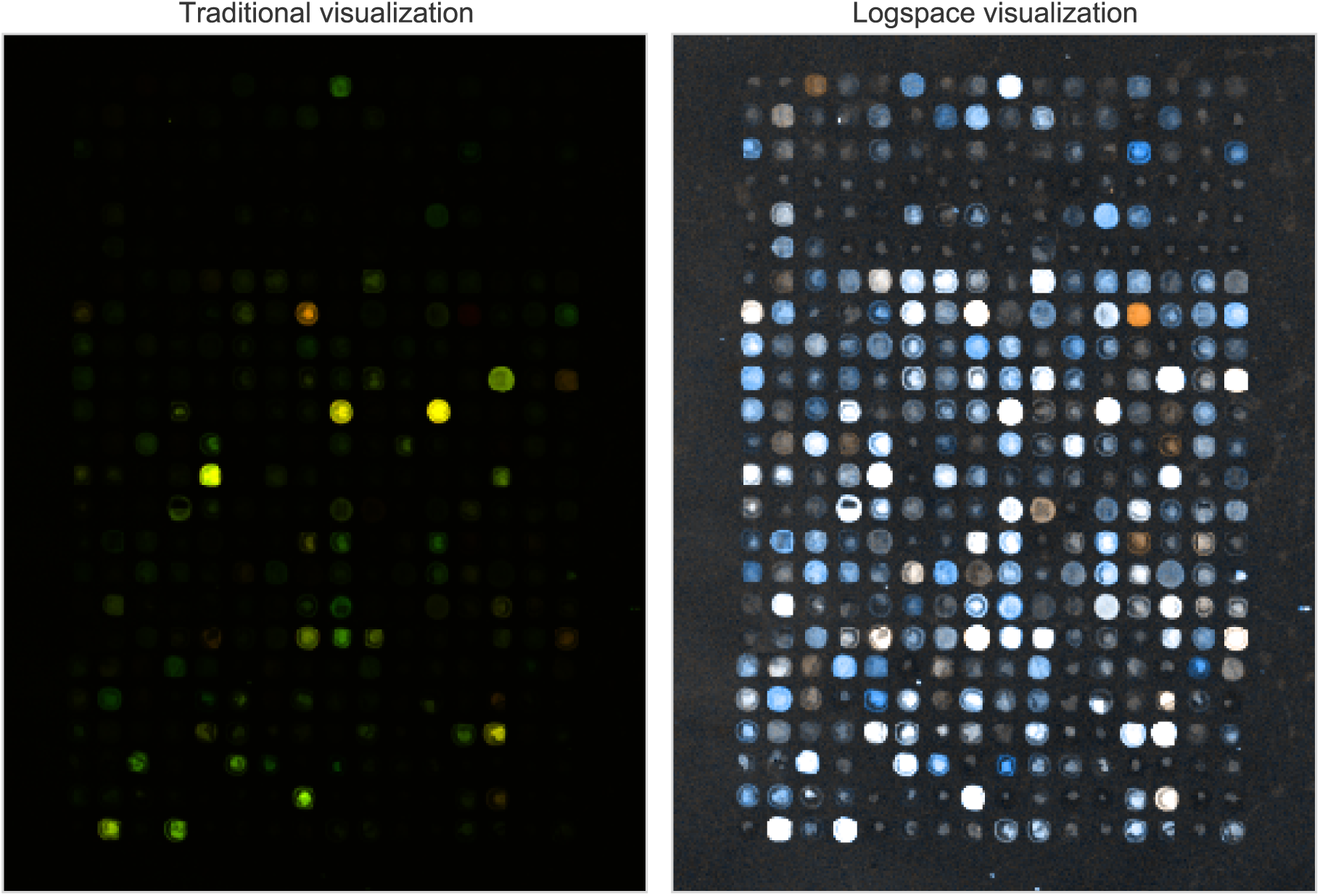
Traditional visualization of a two-channel microarray image versus the log-space visualization.

The problem with this approach is that raw intensity values have a range of [0,65536] (or 216), yet 90% of pixels in a typical microarray have a value below 1,000. Although this representation is perceptually faithful, it can hide many weak spots that are in fact perfectly recognisable, as demonstrated in Figure 2.

As an alternative, we propose mapping channels to 24bit RGB using

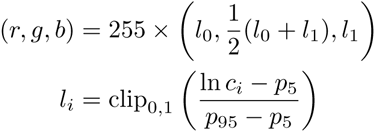

where *c*_*i*_ is the intensity in channel *i* and *p*_5_, *p*_95_ are the fifth and ninety-fifth percentiles of the log-intensities across all channels. The effect of this transformation can be seen in Figure 2.

In the above, *p*_5_ and *p*_95_ were chosen arbitrarily if a specific visualization comes out too light, *p*_5_ can be replaced with a higher percentile, while if the visualization comes out too dark, *p*_95_ can be replaced with a lower percentile.

Normalization aside, we experimented with several different mappings of microarray channels onto colour channels. We found the most aesthetically pleasing to be mapping each channel onto R and B directly, with G being the mean of the two.

## 3 Results

As discussed in Section 2.1, we benchmarked our approach on the two datasets of Lehmussola et al.

### 3.1 Simulated Data Benchmark

On the simulated data benchmark, we tested our model on the 25 low-quality images that were not used for training, along with the 50 high-quality images that were not used for training. As in Lehmussola et al., in Table 2 we report the pixel classification error rate and the discrepancy distance the average distance of a misclassified pixel from the nearest pixel of the correct class. Both metrics were calculated for each image individually, with the medians being reported here.

We provide six algorithms for comparison:

**FC** is the fixed-circle approach, which applies a circular mask of constant radius to each spot.

**KM** is the *k*-means approach of Bozinov and Rahnenführer [3], which forms a foreground cluster and a background cluster from the pixels around a spot based on their intensities in each channel.

**HKM** is the hybrid *k*-means approach also of Bozinov and Rahnenführer, which augments the KM approach with a mask that can be used to eliminate outliers [3].

Together, FC, KM and HKM represent the best-performing algorithms of those reported on in Lehmussola et al. The next three algorithms are drawn from other papers that have reported on the Lehmussola et al. benchmark.

**AG** is the adaptive graph method of Karimi et al. [13] which uses minimum graph cuts to segment the region around a spot into foreground and background.

**3D** is the 3D spot modelling algorithm of Zacharia and Maroulis [21] which models each spot as a 3D model, then tries to approximate the model using a smooth function generated by a genetic algorithm.

**PASS** is the probabilistically-assisted spot segmentation algorithm of Gjerstad et al. [7], which iteratively fits a Gaussian to a spot and uses the fit to decide which pixels should be used to fit the next iteration.

The results of these six plus NN, our neural network method on the simulated data benchmark can be seen in Table 2. Our method matches the best existing methods on three of the four metrics, only falling short on pixel error rate on the low-quality data.

Most of our neural network’s 0.8% error rate on the lowquality data is due to spots similar to the one shown in the lower row of Figure 3. There, the spot is both shifted from the grid and has low expression levels in both channels, making the dislocation invisible even to a human but not to PASS’s statistical approach.

**Figure 3:**
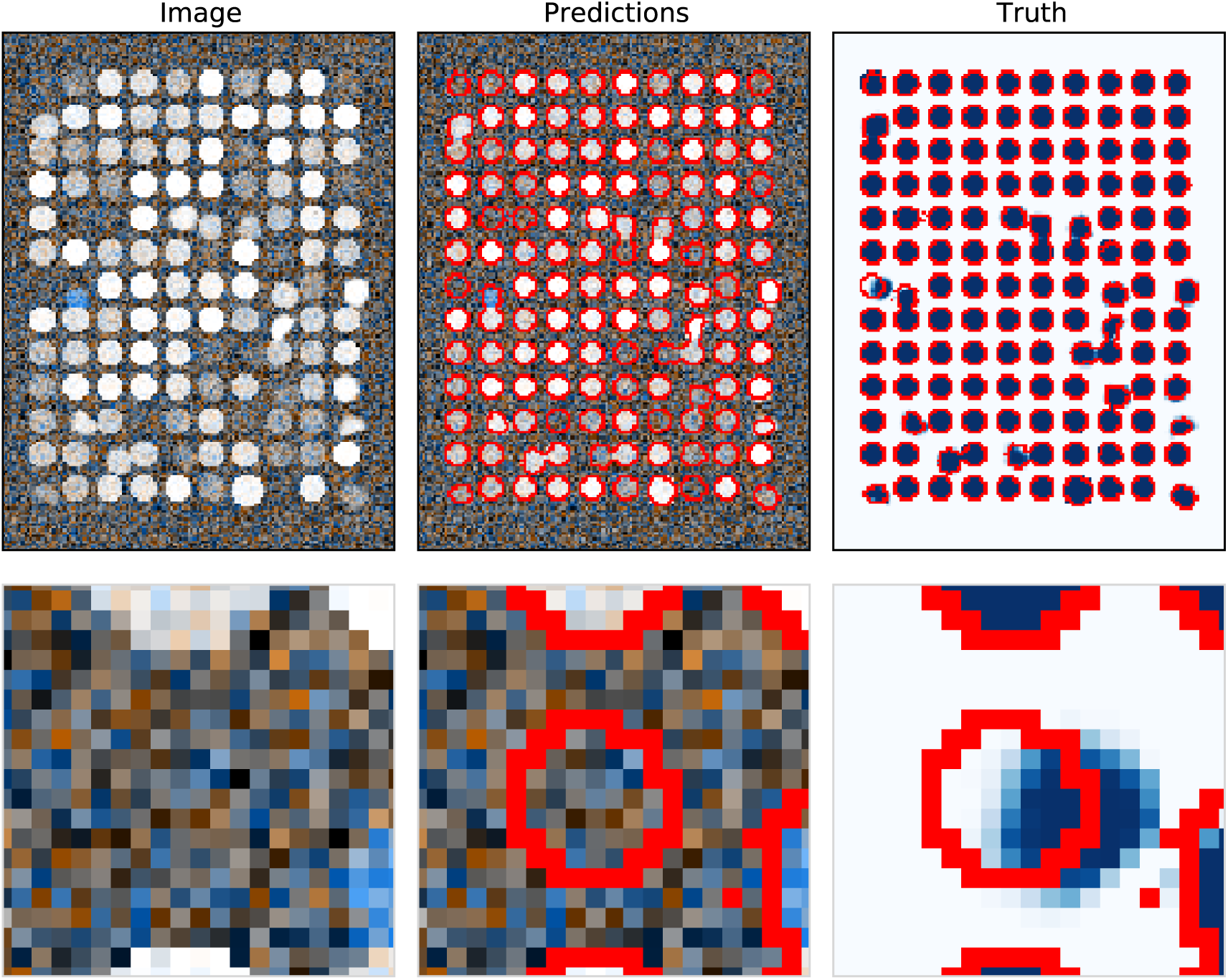
Upper row: example of results for poor-quality simulated data. Lower row: zoom on the spot in seventh row, first column, which has been badly misclassified.

**Table 2:** Performance on the simulated data benchmark.

| Measure | Error (%) |  | Discrepancy (0.01px) |  |
| --- | --- | --- | --- | --- |
| Dataset | Good | Low | Good | Low |
| FC | 4.9 | 4.9 | 2.7 | 2.7 |
| KM | **0.0** | 2.5 | **0.0** | 4.1 |
| HKM | 1.7 | 2.0 | 1.6 | 2.9 |
| AG | 1.2 | 2.1 | 1.7 | 2.9 |
| 3D | **0.0** | 1.2 | **0.0** | 1.8 |
| PASS | **0.0** | **0.0** | **0.0** | **0.0** |
| NN | **0.009** | 0.847 | **0.001** | **0.062** |

### 3.2 Experimental Data Benchmark

On the experimental data benchmark, we tested our model on the four images not used for training. We used a simple Hough-line based gridding heuristic to index the spots identified by our neural network, then calculated the expression ratio of each spot using Lehmussola et al.’s methodology. Their methodology estimates the expression level of a spot as the median of the pixel intensities assigned to the spot, applies a morphological background correction [20] to the levels, then takes the ratio of the corrected levels to be the expression ratio of the spot.

Similarly to Lehmussola et al., we found that about 2,000 of the 12,000 spots were missing in one or more of the images. On the basis of the expression levels of the remaining 10,000 spots, we calculated the six pairwise correlations and mean-absolute-errors, which can be seen in Figure 4. Unfortunately of the six algorithms given as comparison on the simulated data, only the three from Lehmussola et al. have readily-available results on this dataset; the other three all benchmark on their own experimental microarray datasets.

**Figure 4:**
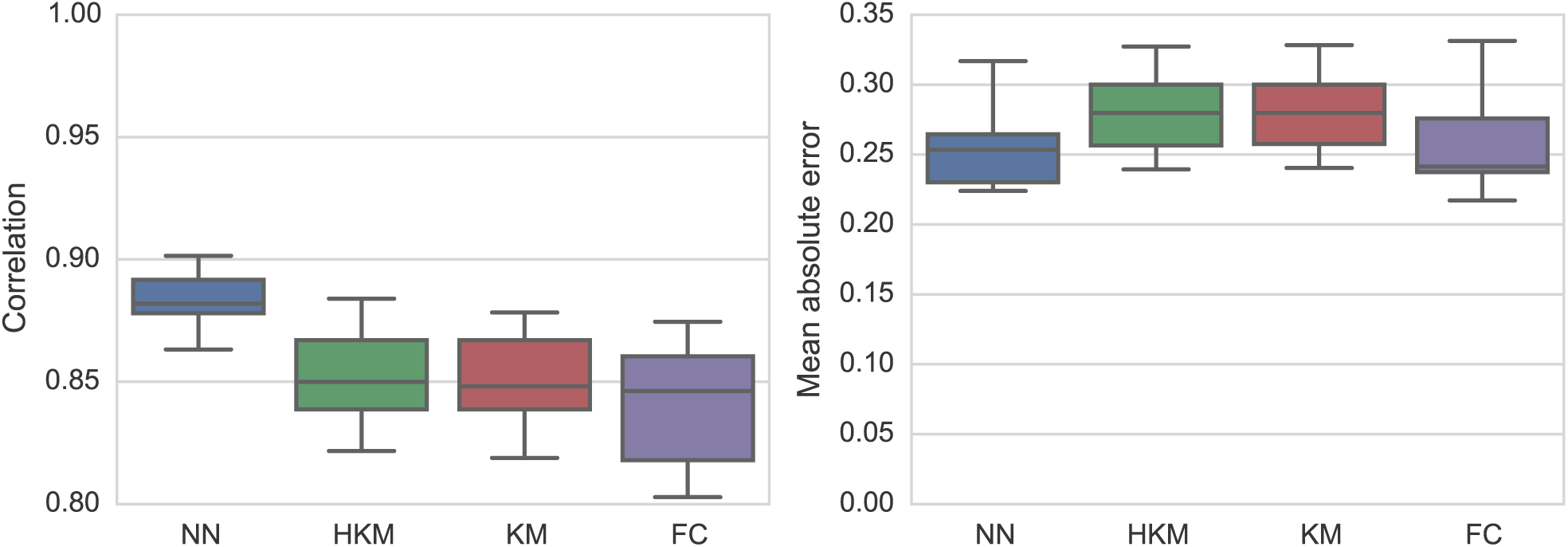
Performance on the experimental benchmark

Our method improves on those detailed in Lehmussola et al. in terms of correlation, but is slightly poorer than the FC method in terms of mean absolute error. We believe this is because the error is in terms of the ratio, which is very sensitive to anomalously small denominators something that is more of a problem for our method, which is happy to assign a spot with an area of just a few pixels. Clipping the expression ratios at the 99th percentile is an easy way to counter this, and leads to a more respectable 0.22 for the median of the mean absolute errors.

An example of the outlines generated by our approach can be seen in Figure 5.

**Figure 5:**
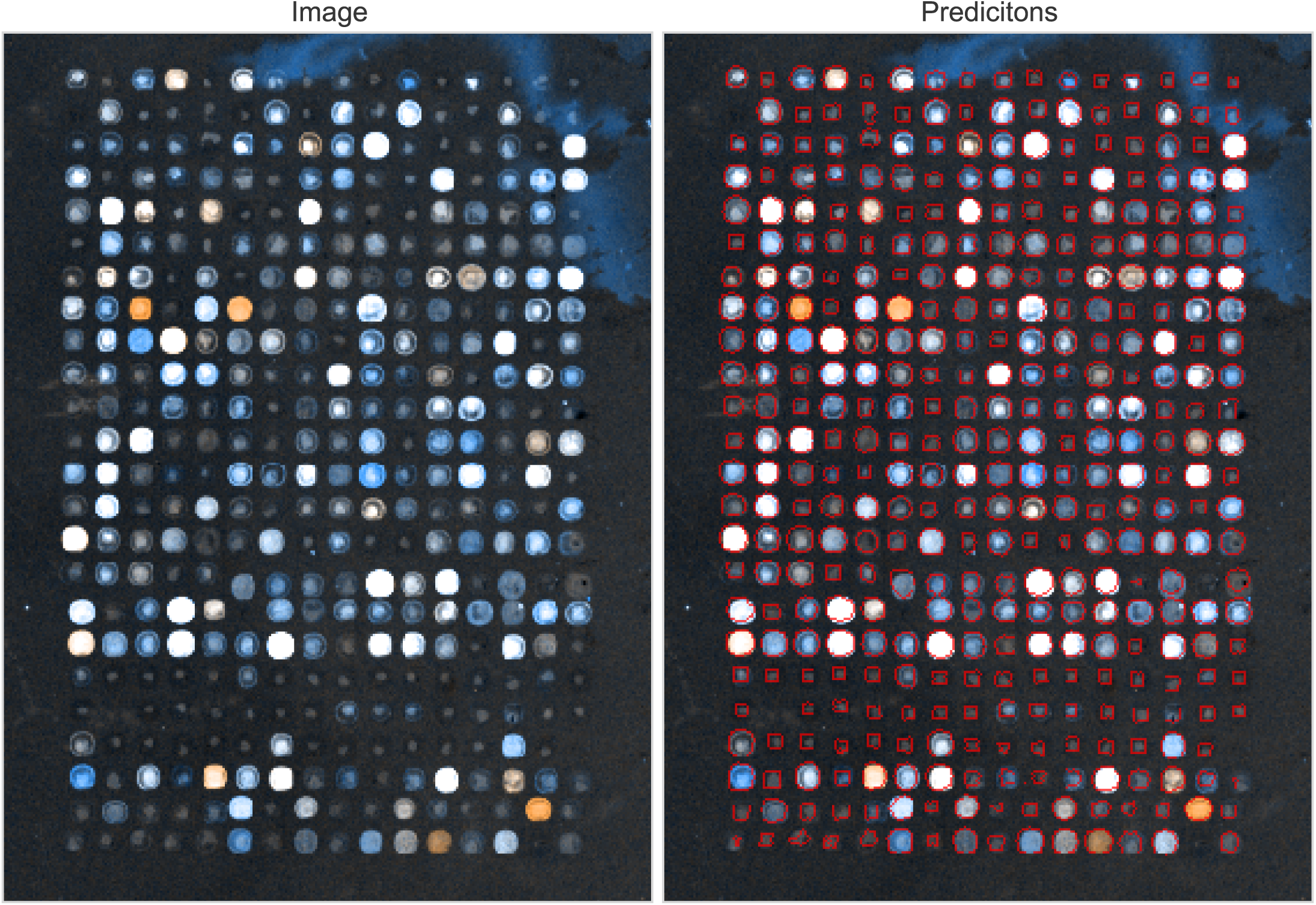
Example of results for experimental data

## 4 Discussion

While our method performs well on the traditional benchmarks, its greatest advantage is in it’s robustness. Array images like Figure 1 are common in experimental work, yet are absent from the microarray segmentation literature.

It’s also notable that our approach and network architecture are both similar to the one used by Ciresan et al. to solve a completely different image segmentation task. This indicates that our approach is likely to generalize very well to different kinds of array and different kinds of corruption. If the network repeatedly misclassifies a certain kind of corrupted spot, all that is required to fix it is for a human to label a few examples and add them to the training set.

Finally, these results were achieved on the back of a fairly simple neural network architecture. As the field matures, we expect the performance of neural networks on these kinds of segmentation problems will improve with it. To encourage this, we have made all our code available on Github1 under a BSD license.

## Footnotes

1 https://github.com/andyljones/NeuralNetworkMicroarraySegmentation

